# Fatigue during acute systemic inflammation is associated with reduced mental effort expenditure while task accuracy is preserved

**DOI:** 10.1101/2023.05.19.541125

**Authors:** B.I.H.M. Lambregts, E. Vassena, A. Jansen, D.E. Stremmelaar, P. Pickkers, M. Kox, E. Aarts, M.E. van der Schaaf

## Abstract

**Background:** Earlier work within the physical domain showed that acute inflammation changes motivational prioritization and effort allocation rather than physical abilities. It is currently unclear whether a similar motivational framework accounts for the mental fatigue and cognitive symptoms of acute sickness. Accordingly, this study aimed to assess the relationship between fatigue, cytokines and mental effort-based decision making during acute systemic inflammation.

**Methods:** Eighty-five participants (41 males; 18-30 years (M=23.0, SD=2.4)) performed a mental effort-based decision-making task before, 2 hours after, and 5 hours after intravenous administration of 1 ng/kg bacterial lipopolysaccharide (LPS) to induce systemic inflammation. Plasma concentrations of cytokines (interleukin (IL)-6, IL-8 and tumor necrosis factor (TNF)) and fatigue levels were assessed at similar timepoints. In the task, participants decided whether they wanted to perform (i.e., ‘accepted’) arithmetic calculations of varying difficulty (3 levels: easy, medium, hard) in order to obtain rewards (3 levels: 5, 6 or 7 points). Acceptance rates were analyzed using a binomial generalized estimated equation (GEE) approach with effort, reward and time as independent variables. Arithmetic performance was measured per effort level prior to the decisions and included as a covariate. Associations between acceptance rates, fatigue (self-reported) and cytokine concentrations levels were analyzed using partial correlation analyses.

**Results:** Plasma cytokine concentrations and fatigue were increased at 2 hours post-LPS compared to baseline and 5 hours post-LPS administration. Acceptance rates decreased for medium, but not for easy or hard effort levels at 2 hours post-LPS versus baseline and 5 hours post-LPS administration, irrespective of reward level. This reduction in acceptance rates occurred despite improved accuracy on the arithmetic calculations itself. Reduced acceptance rates for medium effort were associated with increased fatigue, but not with increased cytokines.

**Conclusion:** Fatigue during acute systemic inflammation is associated with alterations in mental effort allocation, similarly as observed previously for physical effort-based choice. Specifically, willingness to exert mental effort depended on effort and not reward information, while task accuracy was preserved. These results extend the motivational account of inflammation to the mental domain and suggest that inflammation may not necessarily affect domain-specific mental abilities, but rather affects domain-general effort-allocation processes.

## INTRODUCTION

Persistent fatigue and cognitive difficulties that arise during and after infection or medical treatments (e.g. cancer-related fatigue, post intensive care syndrome or long-covid) are increasingly recognized societal problems with debilitating effects on daily and occupational functioning (Gilligan, 2015; Skorvanek et al., 2015). These symptoms can be physical, reflecting increased physical exhaustion and lack of energy to perform physical tasks, but also mental, including concentration problems and cognitive difficulties after tasks that require sustained attention, i.e. “brain fog” (Friedman et al., 2010). It has been suggested that persistent fatigue and cognitive difficulties may result from inflammatory effects on brain functioning (Dantzer et al., 2014; Eisenberger et al., 2010; Felger & Miller, 2012; Harrison et al., 2016; Swardfager et al., 2016; Vancassel et al., 2018). However, it is still debated whether inflammation directly affects cognitive functioning, or whether it changes motivational prioritization, i.e the willingness to expend mental effort for reward.

Within the mental domain, it has been suggested that infections or medical treatments such as chemotherapeutic treatment induce neuroinflammation. This neuroinflammation can have direct domain-specific effects on cognitive abilities by affecting neural plasticity, neurogenesis and meyeliniation of neural circuits that support specific cognitive functions (Lacourt et al., 2018; Monje & Iwasaki, 2022; Ortelli et al., 2021; Sleurs et al., 2022). In contrast, within the physical domain, inflammation does not appear to directly affect physical abilities. Instead, it adaptively changes motivational prioritization to direct energy towards internal processes that fight disease and away from external physical activities (Aubert, 1999; Aubert et al., 1997).

This has been further investigated within the physical domain in animal and human studies that assessed the effects of acute inflammation on effort-based decision-making tasks (Boyle et al., 2019; Draper et al., 2018; Lasselin et al., 2017; Nunes et al., 2014; Yohn et al., 2016). More specifically, in the study by Draper et al. (2018), volunteers chose to exert effort for reward or to do nothing and not receive a reward. Effort and reward levels were parametrically modulated to allow the dissociation of effort and reward influences on choice. They found that in healthy volunteers, systemic inflammation (induced by intravenous LPS administration), resulted in decreased physical effort expenditure by selectively increasing the weighting of physical-effort costs, but not reward, in decisions on whether to engage in effortful activities, while physical ability to perform the effort levels remained intact. This selective effect of inflammation on effort but not reward-weighting is in line with neuroimaging studies showing that effort and reward are processed in partly dissociable neural substrates (Lopez-Gamundi et al., 2021). This suggests that inflammation may also differentially affect these dissociable neural processes that underlie physical effort-based decision making.

fMRI studies comparing cognitive and physical effort-based choice found partly overlapping, but also dissociable substrates for processing cognitive versus physical effort (Chong et al., 2017; Schmidt et al., 2012). Accordingly, for the mental domain it is possible that inflammation may affect similar overlapping motivational processes as previously observed for physical effort, but it is also well possible that it directly affects domain-specific processes.

Within the cognitive neuroscience field, motivation and effort-based decision making have repeatedly been associated with fatigue. It is widely acknowledged that increases in fatigue after prolonged active engagement of physical or mental tasks reduces the subsequent motivation to exert effort for reward (Boksem and Tops, 2008; Kok, 2022; Massar et al., 2018; Muller and Apps, 2019). Inflammation is also known to increase fatigue, likely through its effect on dopamine levels (Dantzer et al., 2014), a neurostransmitter that modulates motivational processing in the brain (Kok, 2022; Muller and Apps, 2019; Westbrook et al., 2020). However, the link between inflammation-induced fatigue, behavioural change and cytokine responses has been conflicting (Boyle et al., 2019, 2020; Draper et al., 2018; Lasselin et al., 2020), likely due to small sample sizes of these studies. Moreover, the direct link between *mental* effort-based decision making and inflammation-induced fatigue has not been investigated yet.

This study assessed a mental effort-based decision-making task (Vassena et al., 2019a; Vassena et al., 2019b) in a large group of young healthy volunteers who were intravenously challenged with LPS to assess 1) whether mental performance or motivational choice is altered during LPS-induced systemic inflammation, 2) if the latter, what components of motivational choices are altered (i.e. effort or reward) and 3) whether individual differences in these changes relate to cytokine responses and/or subjective reports of fatigue. We hypothesized that LPS-induced systemic inflammation affects motivational decision-making similarly to what was observed previously for physical effort-based decision-making: i.e., no change in accuracy and selective effects on effort processing, but not on reward processing. In addition, we hypothesized that observed changes are related to cytokine responses and subjective reports of fatigue.

## METHODS

### Study Design, Participants, and Ethics

This study was part of a larger cohort study performed at the department of Intensive Care Medicine of Radboudumc in Nijmegen, the Netherlands, which aimed to identify genomic and transcriptomic biomarkers of interindividual variability in acute systemic inflammation and immune tolerance *in vivo* (Jansen et al., 2022). In this study, we recruited 113 healthy, non-smoking Caucasian participants aged between 18-30 years who had no medical/psychiatric history and did not use prescription drugs (see supplementary table S1 for inclusion and exclusion criteria). All participants underwent two LPS challenges with an interval of one week. Ninety-five participants of this cohort performed the effort-based decision-making task. Participants refrained from eating food (12h) or drinking any kind of beverage containing alcohol or caffeine (24h) before LPS administration. All study procedures were in accordance with the declaration of Helsinki, and approved by the local medical ethics committee (CMO:2018-4983) and participants provided written informed consent. We compensated participants for their participation (400 euro for two 8 hours sessions). This study focused solely on the first LPS challenge.

### General Session Procedures

We performed all LPS-related procedures as described previously (Jansen et al., 2022). We tested three participants simultaneously in one room at the Medium Care Unit of Radboudumc. Sixty to 90 minutes before LPS administration, we placed a radial artery catheter and antebrachial venous cannula to allow serial blood sampling, hemodynamic monitoring, and administration of fluids and LPS, respectively and these stayed in during the remaining of the test-day. In the 45 minutes prior to LPS administration, we administered hydration fluids (2.5% glucose / 0.45% sodium chloride) as a 1.5L prehydration bolus to reduce the risk of vasovagal collapse(van Eijk et al., 2004), and thereafter at a rate of 150 mL/h for the remainder of the experiment. Directly after prehydration, we administered a bodyweight-adjusted bolus dose of 1 ng/kg LPS (*E. Coli*-derived, Type O113, lot no. 94332B1; List Biological Laboratories) intravenously. Behavioural testing took place at three timepoints: 45 minutes before injection (S1), 2 hours post injection (S2) and 5 hours post injection (S3). Timing was based on previous studies showing that sickness symptoms and cytokines peak 1.5-2 hours after LPS administration and are largely normalized 5 post-LPS administration (Kox et al., 2015; Leentjens et al., 2012). At S2, participants were able to perform behavioural tasks while cytokine levels and fatigue are still high (Draper et al., 2018). Participants filled out mood questionnaires each hour after LPS administration. Furthermore, we obtained blood samples at various timepoints prior to and following LPS administration (see below).

Participants performed the effort-based decision making task while sitting in a hospital bed with a laptop computer placed on a table in front of them. To reduce distractions and interference, we separated participants with curtains, gave them noise-reducing headphones during task-performance, and instructed them not to talk to each other about the task or about choice strategies. We also instructed all (medical) personnel in the room not to disturb the participants while they performed the task. The task started approximately 5 minutes past the hour, directly after the hourly sampling of blood, temperature, heartrate and questionnaires.

### Experimental Task

We adjusted the effort-based decision-making task from Vassena et al. (2014, 2019), and presented it on a laptop computer using E-Prime 2.0 software (Psychology Software Tools, Pittsburgh, PA). Participants performed 3 sessions in total (45 minutes before injection (S1), 2 hours post injection (S2) and 5 hours post injection (S3.) Each session included a training phase, a test phase and a choice phase.

During the training phase, participants first performed 21 (S1) or 12 (S2 and S3) example calculations to get familiarized with the easy, medium and hard calculations and task speed. Training was shorter in S2-3 as the participants were already familiar with the task. This was followed by 36 test trials (12 per effort level), during which the task provided feedback (correct/incorrect). No reward was included in these trials. We used these test trials to measure accuracy (proportion correct) per effort level and session.

During the choice phase, the task presented participants with a series of offers in which they decided whether they were willing to accept and perform a calculation of a certain difficulty level to obtain a given reward (See Figure 1A for details on timing). The task presented difficulty and reward levels simultaneously at the beginning of each trial, using the words ‘easy’, ‘medium’ and ‘hard’ to indicate the difficulty level, and the words ‘5 points’, ‘6 points’ and ‘7 points’ to indicate the reward level. We sampled each combination 15 times in a random order, totaling 135 trials per session. Next, the words ‘reject’ and ‘accept’ appeared at the bottom left and right of the screen (location randomized) and participants indicated their choice with their left or right index finger. If participants accepted the offer, the calculation appeared on screen directly after the choice. Calculations were additions and subtractions of 4 single-digit numbers and we manipulated difficulty by the number of carrying and borrowings in the calculation (e.g. easy: 5+6**+1+1**, medium: 5+6-8**+1**, hard: 5+6+9-8), a procedure known to elicit robust differences in difficulty in behavioral performance and willingness to exert effort (Vassena et al., 2019). After the calculation, two possible results appeared on the screen and participants indicated their answer with their left or right index finger. If correct, the screen showed ‘correct’ and the amount of points won (5, 6 or 7 points). If incorrect or too late, the screen showed ‘incorrect’ or ‘too late’ and ‘-1 point’ appeared on screen. If the offer was rejected, participants waited for the duration of the trial and were shown ‘+4 points’ at the end of the trial. Total trial duration was identical for accept and reject trials (7700 ms). After the choice phase, participants rated perceived difficulty (‘how difficult did you find the calculation preceded by these words?’) and enjoyment (‘how enjoyable did you find the calculation preceded by these words?’) of each of the nine offers on a 7-point Likert scale (not at all – very much). This procedure has been used in previous studies to confirm subjective perception of difficult trials as more difficult (Vassena et al., 2014). Task outcome measures for analysis were acceptance (yes/no) reaction times during the choice phase, and accuracy during the training phase.

**Figure 1.**
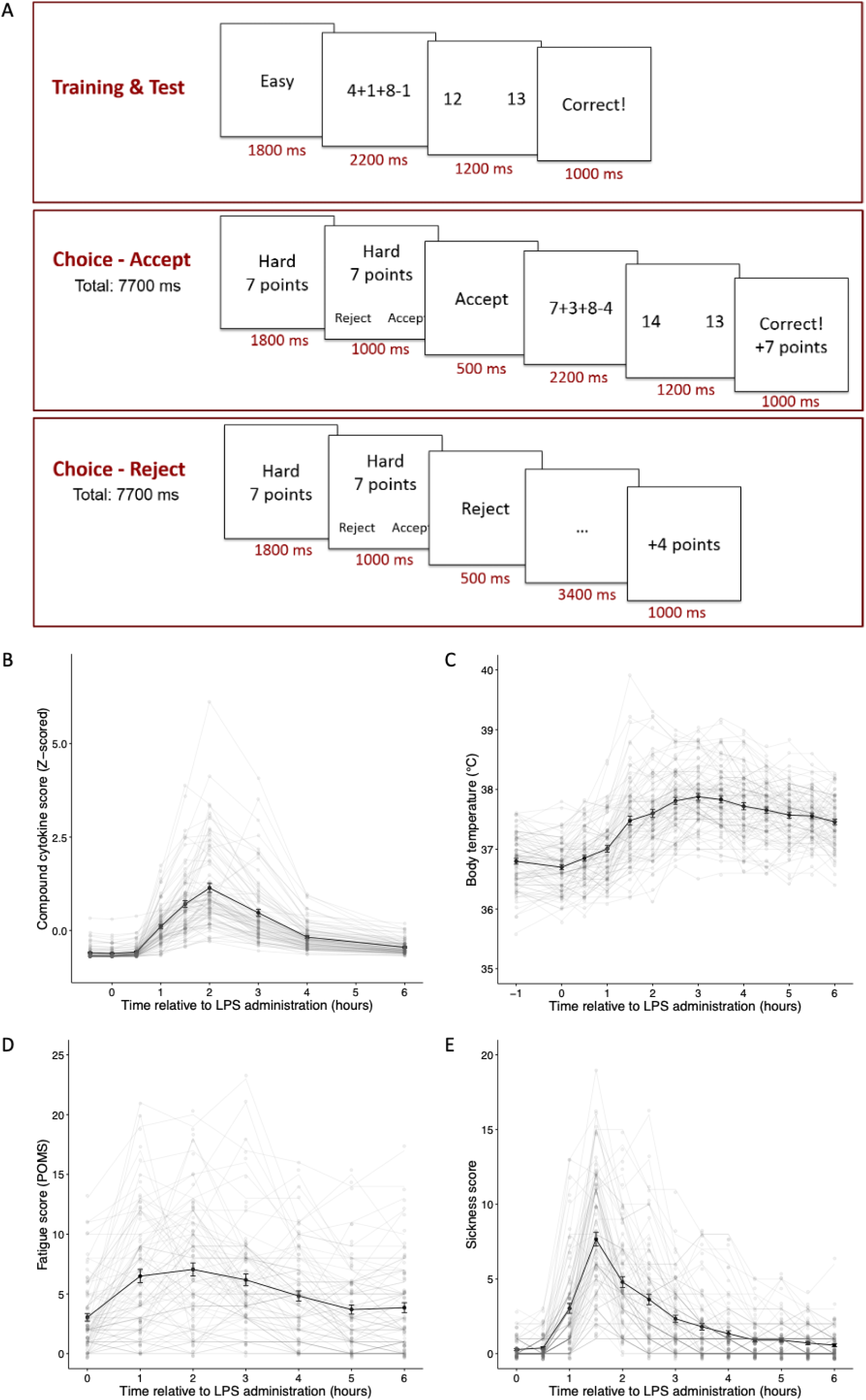
A. Schematic visualisation of the task design and timing. Presented screens per trial for training and test phase (upper panel). Presented screens per trial for the choice phase in case of offer acceptance (middle panel) and offer rejection (lower panel). B. Time course of the composite score of plasma concentrations of cytokines IL-6, IL-8 and TNF. C. Time course of body temperature. D. Time course of state fatigue levels as measured with the subscale fatigue of the Profile of Moods State (POMS) Questionnaire – subscale fatigue scores over time. E. Time course of the sickness scores. Sub-plots B, C, D and E show the mean time course in black with error bars (standard error of the mean) and individual time courses in light grey. IL = Interleukin, TNF = Tumour Necrosis Factor, LPS = Lipopolysaccharide *2-column fitting image*

Participants received a surprise bonus at the end of the study based on the amount of points they had collected (grams of chocolate money).

### Measurements of fatigue, sickness behavior and cytokine levels

An overview of all questionnaires assessed during the session is provided in the supplementary materials (Table S2). For this study, we assessed fatigue using the fatigue subscale of the profile of moods state questionnaires (POMS) (McNair et al., 1989) hourly on iPads using an electronic data capture system (Castor EDC) between S1-S3. We assessed sickness symptoms (headache, muscle pain, back pain, nausea, shivers and vomiting) orally on a 6-point Likert scale every 30 minutes between S1-S3. We determined body temperature every 30 minutes between S1-S3 using an infrared tympanic thermometer (Covidien, Dublin, Ireland). We collected ethylenediaminetetraacetic acid (EDTA)-anticoagulated blood eight times between S1-S3 (–0.5, 30, 60, 90, 120, 180, 240, and 360 minutes relative to LPS administration), immediately centrifuged (2000 g, 4°C, 10 min), after which we stored plasma at –80°C until analysis. We determined concentrations of cytokines of interest (IL-6, IL-8 and tumor necrosis factor (TNF)) using a simultaneous multiplex assay (MILLIPLEX xMAP Human Cytokine/Chemokine Magnetic Bead Panel – Immunology Multiplex Assay, catalogue number HCYTOMAG-60K) in a single batch. We chose to focus on the pro-inflammatory cytokines IL-6, IL-8 and TNF based on previous literature highlighting the importance of these cytokines in various chronic conditions (Bower & Lamkin, 2013; Dowlati et al., 2010; Schubert et al., 2007) and effects of cytokine administration on effort-based decision making in animals (Vichaya et al., 2014; Yohn et al., 2016). Lower detection limits in plasma and intra-assay coefficients of variation (C.V.) were 0.9 pg/ml (C.V. 2.0%) for IL-6; 0.4 pg/ml (C.V. 1.9%) for IL-8; and 0.7 pg/ml (C.V. 1.6%) for TNF.

### Statistical analysis

#### Behavioral Task

We analyzed acceptance data (yes/no) using a generalized estimated equation (GEE) with a binary logistic model and independent correlation structure (Treadway et al., (2009). We chose GEE because it is particularly suitable for repeated measures designs of non-normal binary data, and allows to specify different correlation structures for different timepoints and conditions within one group/subject (Hubbard et al., 2010). Acceptance was entered in the model as dependent variable with Effort level, Reward level and Session as factors. To control for learning across sessions, individual accuracy levels were added, calculated as the proportion correct per effort level during the training phase, as a covariate to the model. A GEE model with an independent correlation structure resulted in the best fit (QIC = 24996) compared to an exchangeable (QIC=918642) or ar1 (QIC=25160) correlation structure.

First, we evaluated whether participants showed the expected behavior: main effects of effort and reward across all three sessions (i.e. higher acceptance rates for offers with higher rewards and lower effort). Next, we assessed the whether there was an effort by session, reward by session or effort by reward by session interaction, including all three sessions. As in Draper et al. (2018), we then assessed whether acceptance rates decreased between S1-S2. If this analysis revealed a significant result, we tested whether this change normalized at S3 (recovery) by comparing S2-S3 and S1-S3. To assess whether mental accuracy was affected by LPS, we assessed changes in accuracy separately using repeated measures ANOVA with the factors Effort, Reward and Session.

We square-root transformed and analyzed response times (RTs) on choice (moment of accept/reject) using a GEE with a Gaussian model and independent correlation structure. We entered RTs as dependent variable with Effort level, Reward level and Session as factors and accuracy as continuous covariate. Finally, to validate the task manipulation, we conducted two repeated measured ANOVAs with difficulty and enjoyment ratings as dependent variables and Effort, Reward and Session as independent variables.

#### Physiological and subjective responses to LPS

We assessed changes in body temperature, fatigue and sickness scores using the three measurements that we took prior to each task session (i.e. S1, S2 and S3). As we did not perform blood sampling at S3, we calculated cytokine datapoints for S3 by interpolating T=4 hours and T=6 hours.

We baseline-corrected all cytokine datapoints by subtracting their individual plasma concentrations measured 45 minutes pre-LPS administration. We calculated a composite measurement for cytokines by averaging the baseline-corrected and Z-scored individual plasma concentrations of IL-6, IL-8 and TNF for each participant at each timepoint. We calculated fatigue scores as mean score on the POMS subscales for each session. We determined a composite sickness score by adding the scores on all symptoms except vomiting, resulting in a total score between 0-25. We excluded vomiting from the total sickness score as it occurred only in a few subjects.

We conducted our repeated-measures one-way ANOVAs to assess whether cytokine levels, body temperature, fatigue and sickness scores were increased at S2 compared to S1 and S3 with composite cytokine score, body temperature, fatigue score or composite sickness score as dependent variable and Session (3 levels) as independent variable, followed up by repeated-measures ANOVAs comparing two sessions.

#### Relationship between Choice behavior, Fatigue and Cytokines

We performed partial correlation analyses to assess relationships between changes in choice behavior with changes in fatigue and cytokines (composite score). We focused the analysis on the task condition(s) that showed a significant change at S2. Similar to our behavioral analyses, we first compared S1-S2, followed by S2-S3 if S1-S2 proved significant. To this end, we calculated difference scores (e.g. S2 – S1) for the proportion accepted offers (Δacceptance), fatigue (Δfatigue) and the cytokine composite score (Δcytokines). We performed two separate partial correlations analysis: one with Δacceptance as dependent variable, Δfatigue as independent variable and Δaccuracy as controlling variable and one with Δacceptance as dependent variable, Δcytokines as independent variable and Δaccuracy as controlling variable. To control for potential variance related to sex, we repeated these partial correlations with sex as additional controlling variable. We also explored alternative calculations of cytokines: peak IL-6 and a composite score for the area under the curve (AUC) of IL-6, IL-8 and TNF. To this end, we Z-scored and averaged each cytokines’ baseline-corrected AUC for each participant.

Lastly, we assessed the relationship between Δfatigue and Δcytokines by performing a correlation analysis. All assumptions for correlation analyses were met. We defined outliers as a value higher than mean +3SD on the difference scores and excluded those from analyses.

We performed all analyses using R (version 4.0.2) (R Core team (2020), R Foundation for Statistical Computing, Vienna, Austria). We performed main analyses using R packages ‘geepack’, ‘car’ and ‘ppcor’.

## RESULTS

### Participants

We excluded ten participants from analysis because of insufficient proficiency of the Dutch language (N=3), technical failures (N=3), a too low dose of LPS administered (N=3) and incorrect task comprehension (N=1). We did not exclude subjects who vomit, as this did not interfere with task performance. In total, we included 85 participants (41 males, mean age=23.0 years (SD=2.4)) for the final analysis. As one participant did not fill out the difficulty and enjoy ratings, we performed repeated measures ANOVAs on difficulty and enjoy with 84 participants. Questionnaire (i.e. fatigue) data was missing from two additional participants. Accordingly, we performed regression analyses including fatigue data with 83 participants.

### Systemic inflammation, body temperature, fatigue and sickness score increased after LPS administration

Repeated-measures ANOVA revealed a main effect of Session for circulating concentrations of pro-inflammatory cytokines (i.e. the composite measure, reflecting the extent of the systemic inflammatory response) (F_2,84_ = 172.84, p = <0.001). Cytokine concentrations increased on S2 compared to S1 (S1-S2: F_1,85_ = 209.56, p = <0.001). This effect only partly recovered at S3 (S2-S3: F_1,85_ = 140.44, p = <0.001; S1-S3: F_1,85_ = 90.80, p = <0.001) (Figure 1B).

We observed a main effect of Session for body temperature (F_2,84_ = 85.41, p = <0.001). Body temperature increased on S2 compared to S1 (F_1,85_ = 121.7, p = <0.001). This effect did not recover at S3 (S2-S3: F_1,85_ = 0.124, p = 0.725; S1-S3: F_1,85_ = 170.08, p = <0.001) (Figure 1C).

We observed a main effect of Session for fatigue (F_2,84_ = 25.23, p = <0.001). State fatigue increased on S2 compared to S1 (F_1,83_= 40.16, p = <0.001) and this effect fully recovered at S3 (S2-S3: F_1,83_= 25.12, p = <0.001; S1-S3: F_1,83_= 1.69, p = 0.196) (Figure 1D).

For sickness scores, repeated-measures ANOVA revealed a main effect of Session for S1-S2 (F_1,85_ = 150.41, p = <0.001). Self-reported sickness symptoms increased on S2 compared to S1 and this effect was partly recovered at S3 (S2-S3: F_1,85_ = 102.25, p = <0.001: S1-S3: F_1,85_ = 18.21, p = <0.001) (Figure 1E).

### Difficulty and enjoyment ratings matched the intended task manipulation

Participants experienced the effort and reward levels as expected (Figure 2). Repeated-measures ANOVA revealed a main effect of Effort (F_2,84_ = 847.05, p = <0.001) (Figure 2A), but not Reward (F_2,84_ = 1.05, p = 0.350) (Figure 2B) on difficulty ratings. Participants perceived hard trials as more difficult than medium trials (T_84_= 351.35, p = <0.001) and medium trials as more difficult than easy trials (T_84_= 525.68, p =<0.001). Difficulty ratings did not decrease with Session (F_1,84_= 0.63, p = 0.426).

**Figure 2.**
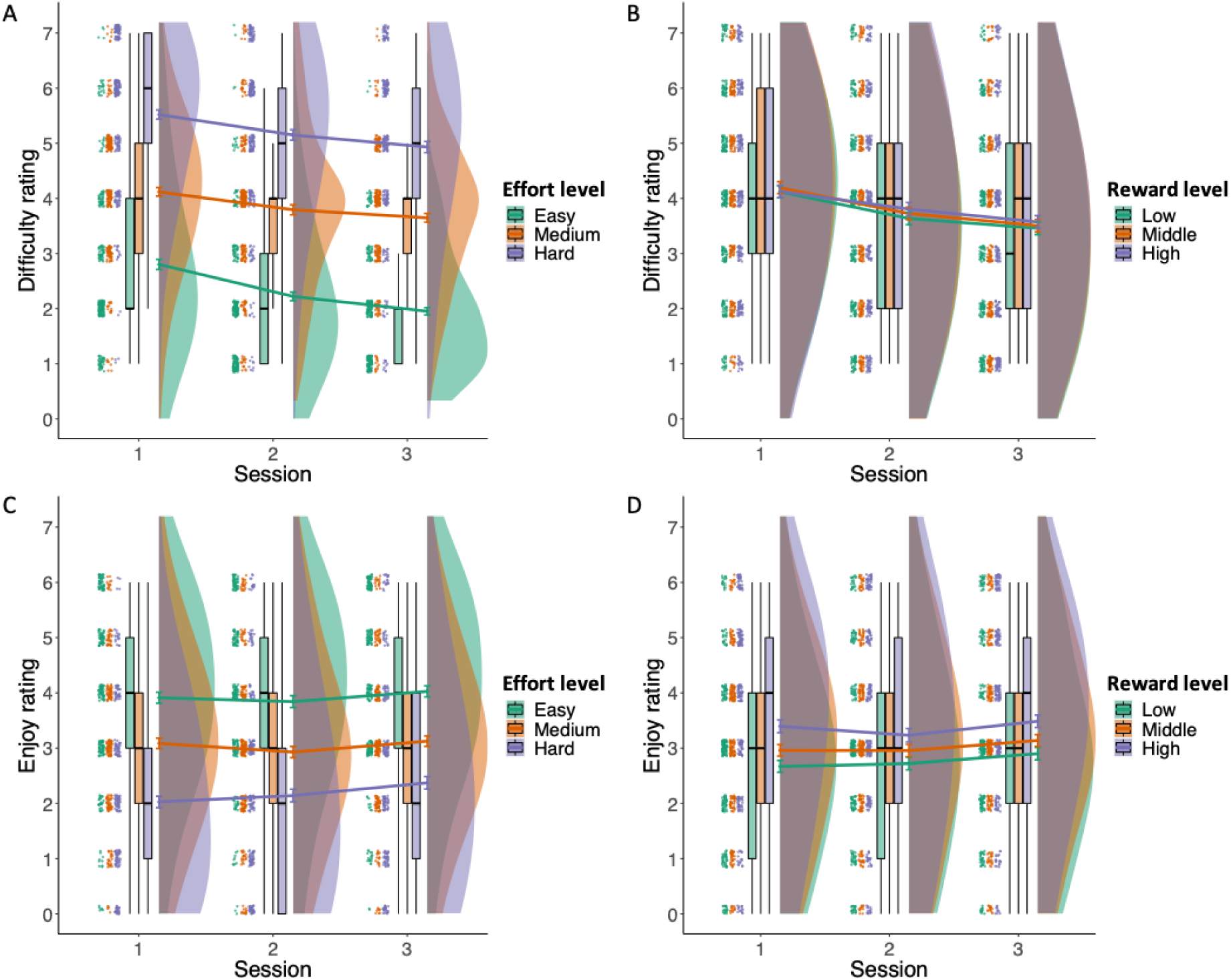
Self-reported ratings for perceived difficulty for the effort (A) and reward levels (B) and enjoyment ratings for effort levels (C) and reward levels (D) across sessions. Each sub-figure shows the individual datapoints jittered around the questionnaire score (7-point Likert scales), box plots with median and standard deviation and violin plots showing the data distribution. The lines show the means with standard error of the mean (SEM). *2-column fitting image*

Repeated-measures ANOVA revealed a main effect of Effort (F_2,84_ = 217.18, p = <0.001) (Figure 2C) and Reward (F_1,84_ = 26.61, p = <0.001) (Figure 2D) on enjoyment ratings. Participants enjoyed the trials with a larger reward more (low-middle: T_84_= 9.90, p = 0.002; middle-high: T_84_= 17.34, p = <0.001) and the trials requiring more effort less (easy-medium: T_84_= 117.97, p = <0.001; medium-hard: T_84_= 106.16, p = <0.001). There was no main effect of Session on enjoyment ratings (F_1,84_ = 2.47, p = 0.116). We did not observe significant interaction effects for difficulty or enjoyment ratings.

### Increased effort but not reward sensitivity on session 2

Results on acceptance rates are presented in Figure 3 and Supplementary tables S3-S6. Participants showed the expected decision-making behavior across sessions. Specifically, GEE analysis revealed main effects for both Reward (Chi-square = 133.71, p= <0.001) and Effort (Chi-square = 263.16, p= <0.001) across all three sessions. Participants accepted more offers with higher Reward levels (low-middle: Chi-square = 84.71, p = <0.001, middle-high: Chi-square = 34.30, p = <0.001) and accepted less offers with higher Effort levels (easy-medium: Chi-square = 94.25, p = <0.001; medium-hard: Chi-square = 98.25, p = <0.001).

**Figure 3.**
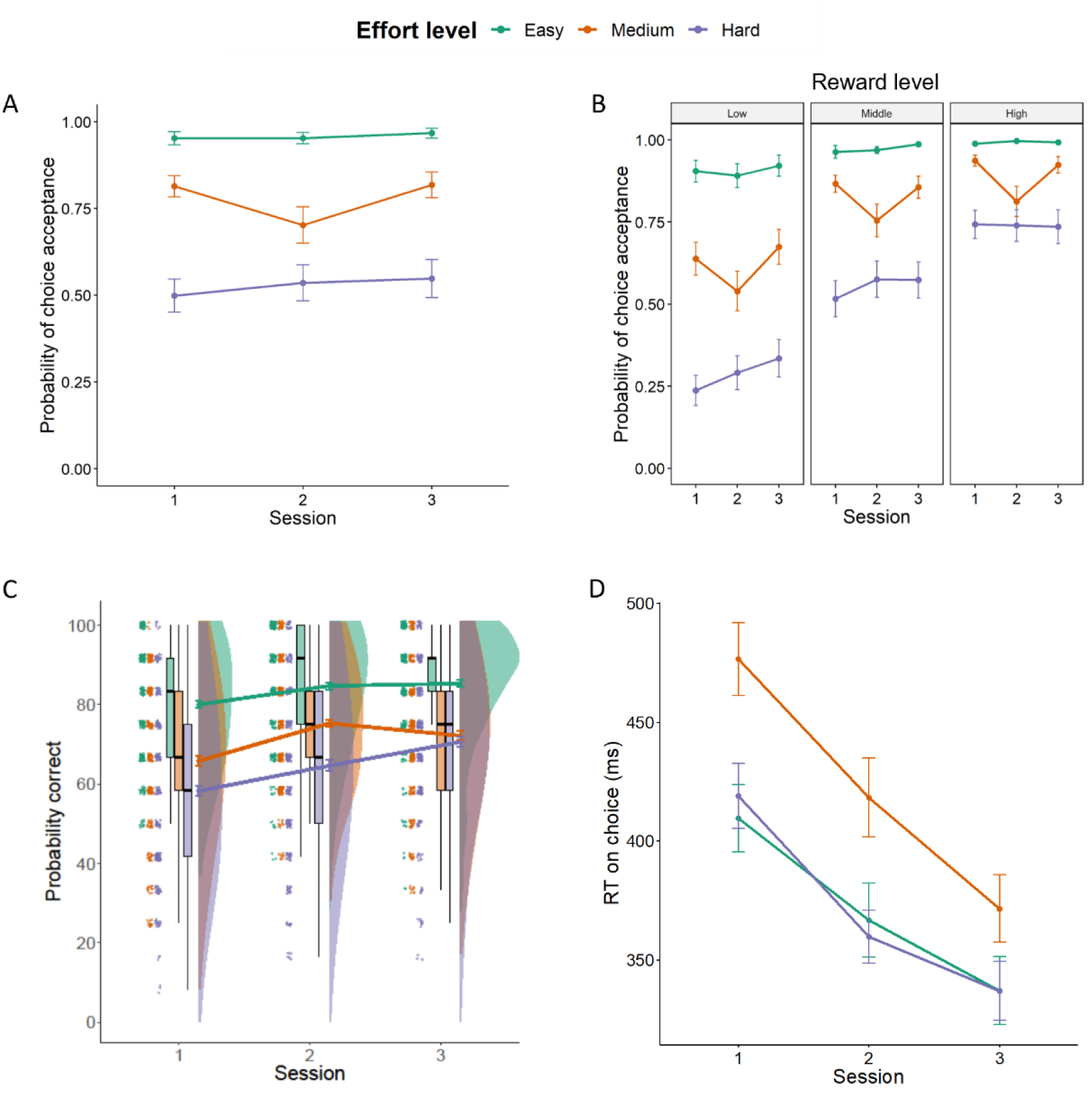
A. Estimated means from the GEE model for choice acceptance split for effort level for all three sessions. B. Estimated means from general estimated equations (GEE) model for choice acceptance split for effort level (color) and reward level (low, medium, high) for all three sessions. C. Accuracy scores based on the test phase of the experiment per session for each participant. Accuracy scores were calculated as the percentage of correct answers for each effort level separately. D. Estimated means from GEE model for response times (RTs) split for effort level for all three sessions. All error bars are standard error of the mean (SEM). For visualization of the uncorrected raw acceptance data and individual trajectories, see supplement figure S2. *2-column fitting image*

When comparing the choice data from all three sessions, the GEE revealed a significant Effort*Session interaction (Chi-square = 17.02, p= 0.002) (Figure 3A), but no Reward*Session interaction (Chi-square = 8.15, p= 0.086) or Effort*Reward*Session interaction (Chi-square = 13.55, p= 0.094) (Figure 3B).

When comparing between S1-S2, the GEE revealed a significant Effort*Session interaction (Chi-square = 14.77, p = <0.001). Breakdown of this interaction by Effort level revealed that at S1-S2, participants accepted less offers with a medium effort level (Chi-square = 9.25, p = 0.002), which was not the case for easy or hard effort levels (easy: Chi-square = 1.81, p = 0.179; hard: Chi-square = 0.30, p = 0.581) (figure 2C). We did not observe a Reward*Session interaction (Chi-square = 0.200, p = 0.905) or Effort*Reward*Session interaction (Chi-square = 8.80, p = 0.066). Thus, the reduction in acceptance rates on medium effort trials did not differ between reward levels (figure 2B).

The acceptance rates for medium effort trials fully restored at S3. The GEE analysis comparing S2-S3 showed an Effort*Session interaction (Chi-square = 9.05, p = 0.011) and no Reward*Session interaction (Chi-square = 2.43, p = 0.296). Participants accepted more offers with a medium effort level again for S3 compared to S2(Chi-square = 17.61, p = <0.001), while we did not observe a change in acceptance rates for offers with easy and hard effort levels (easy: Chi-square = 0.08, p = 0.776; hard: Chi-square = 0.12, p = 0.731). We confirmed the restoration of acceptance rates for medium effort by a lack of Effort*Session (or Reward*Session) interaction when comparing S1-S3 (Effort*Session: Chi-square = 1.97, p = 0.373, Reward*Session: Chi-square = 0.79, p = 0.673).

Importantly, the reduction in acceptance rates for effort level medium from S1-S2 was not reflected in a deterioration in arithmetic accuracy, as participants’ accuracy improved across sessions on all three effort levels (Figure 3C). Repeated-measures ANOVA revealed main effects of Session (F_1,85_ = 54.34, p = <0.001) and Effort (F_1,85_ = 169.44, p = <0.001) and no Effort*Session interaction (F_2,85_ = 2.29, p = 0.10) for S1-S2. For S2-S3, we observed a main effect of Effort (F_2,85_ = 129.67, p = <0.001) and a significant Effort*Session interaction (F_2,85_ = 8.74, p = <0.001), where accuracy only increased for hard trials versus easy (T_85_ = 242.93, p = <0.001) and medium (T_85_ = 27.04, p = <0.001) trials.

### Decision response times for medium effort level were consistently higher across sessions

GEE analysis on decision RT data (acceptance) revealed significant main effects of Effort (Chi-square = 178.60, p = <0.001) and Session (Chi-square = 29.68, p = <0.001). Participants were slower on medium effort trials, compared to easy and difficult trials (easy-medium: Chi-square = 50.67, p = <0.001, easy-hard: Chi-square = 0.42, p = 0.516, medium-hard: Chi-square = 96.26, p = <0.001) and participants became faster with each session (S1-S2: Chi-square = 9.20, p = 0.002, S2-S3: Chi-square = 13.05, p = <0.001) (Figure 3D).

### Relationship between changes in acceptance rates, fatigue and the systemic inflammatory response

One participant was an outlier with respect to their S2-S1 acceptance difference score and three participants were outliers with respect to their S3-S2 acceptance difference score. Additionally, one participant was an outlier with respect to their S2-S1 composite cytokine difference score. We therefore excluded these participants from the partial correlation analyses.

As results showed session effects on medium effort levels, partial correlation analyses on S2-S1 focused on the relationships between Δcytokines, Δfatigue and Δacceptance of medium effort trials only. This revealed a significant negative relationship between Δfatigue and Δacceptance (r = –0.024, p = 0.035) (Figure 4A). Larger increases in fatigue were associated with larger decreases in acceptance rates for medium effort trials. However, restoration of fatigue on S3-S2 was not associated with restoration of acceptance rates (r = 0.06, p = 0.553) (Figure 4B).

**Figure 4.**
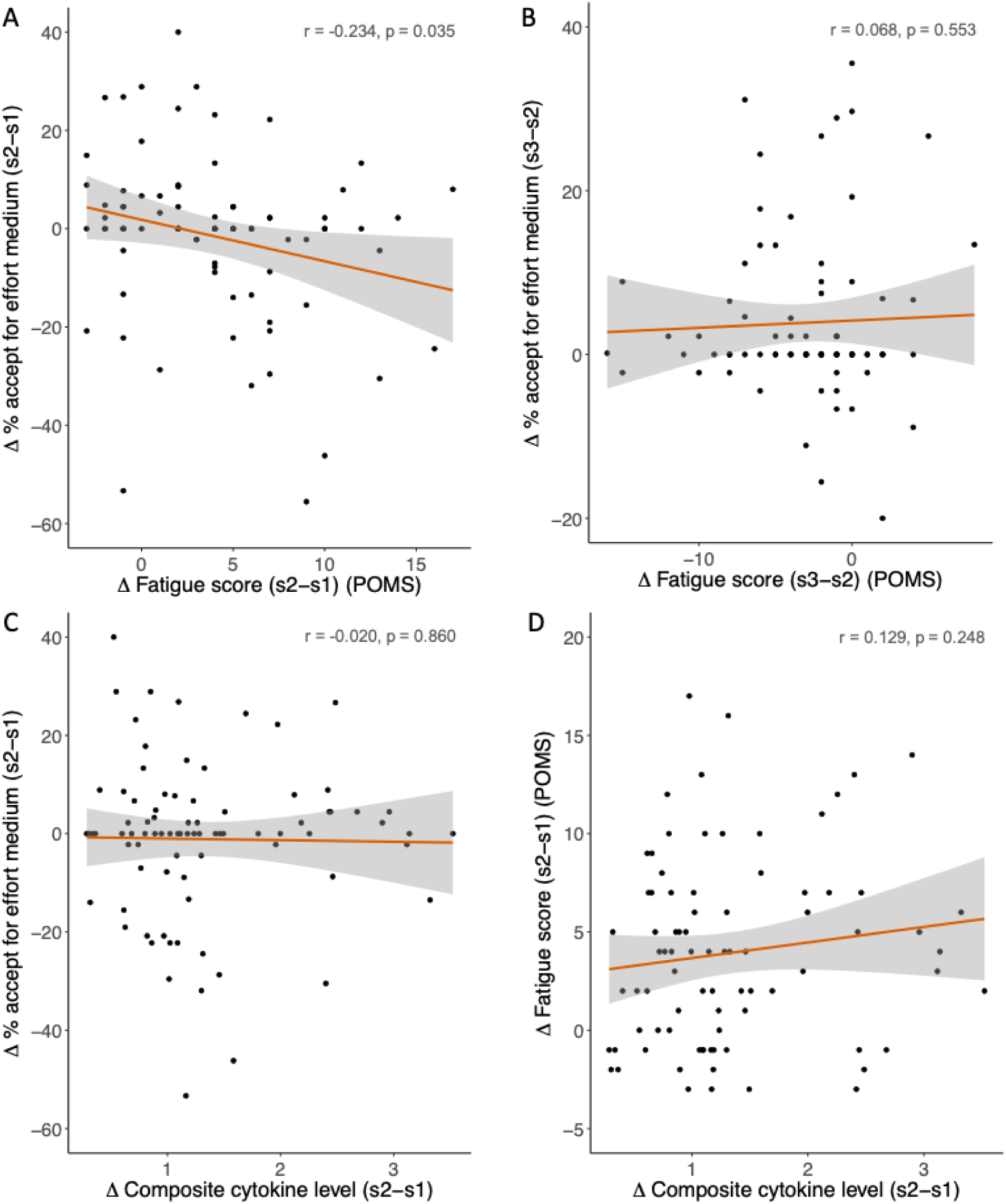
Associations between Δacceptance on medium effort trials and Δfatigue using the subscale fatigue of the Profile of Moods State (POMS) questionnaire for S2–S1 (A) and S3–S2 (B). Panel C shows the association for Δacceptance and the Δcomposite score for peak levels of interleukin(IL)-6, IL-8 and Tumor Necrosis Factor (TNF) for S2–1. Panel D shows the association between Δcomposite cytokine score and Δfatigue for S2–S1. For visualization of correlations at S1 and S2 see supplementary figure S3.

In contrast, we did not observe an association between Δcytokines and Δacceptance on S2-S1 (r = –0.02, p = 0.860) (Figure 4C). Exploration of alternative calculations of cytokines also did not reveal any relationship between Δcytokines and Δacceptance. These explorations included using the area under the curve of the composite cytokine score (r = –0.02, p = 0.856) and peak IL-6 value for S2-S1 (r = – 0.11, p = 0.309). In addition, adding sex as a covariate to the model did not change the result (r = –0.03, p = 0.203). Finally, we did not observe a relationship between Δfatigue and Δcytokines (r = 0.13, p = 0.248) on S2-S1 (Figure 4D).

## DISCUSSION

This is the first study to date that investigated the impact of acute systemic inflammation on mental effort-allocation and task accuracy using the experimental human endotoxemia model, a highly standardized and reproducibe model using LPS as the inflammatory stimulus. The results add to previous studies in several ways. First, they show that systemic inflammation affects mental effort-allocation, despite improved accuracy on arithmetic calculations. Second, systemic inflammation affected mental effort-based choice similarly as observed previously for physical effort-based choice; changes in mental effort-based choice after LPS administration depended on effort and not reward information (Draper et al., 2018). Third, the results show that the observed changes in mental effort-based choice were best predicted by subjective self-reported fatigue, rather than by the extent of the cytokine response.

This study has shown that acute systemic inflammation reduced the willingness to exert mental effort, and that this effect was driven by difficulty of the mental task and not by rewards that could be gained. Acute systemic inflammation did not impair arithmetic performance, as accuracy measurements assessed during the training session before each choice phase revealed improved rather than reduced accuracy over sessions. This finding is consistent with reports on physical effort expenditure where participants were still able to perform the effort levels adequately (Dantzer, 2001; Draper et al., 2018) as well as with previous studies reporting no change in general cognitive functioning after LPS administration (Cohen et al., 2003; Grigoleit et al., 2010; Van den Boogaard et al., 2010). Together, these results support the motivational account of inflammation (Dantzer, 2001) in the mental domain and show that acute systemic inflammation does not impair cognitive performance or mental capacity per se, but rather changes motivational prioritization by reducing the willingness to invest mental effort for reward.

LPS-induced systemic inflammation affected the impact of effort information on choice, but – contrary to previous physical effort-based decision-making tasks that found larger reductions for higher effort levels – acceptance rates were only reduced for medium and not for easy or hard trials. One explanation for this non-linear effect is that LPS-induced inflammation affects choices based on the expected benefit of invested effort, rather than the difficulty of the calculation. Indeed, for easy calculations, high accuracy (approaching ceiling level) was achieved relatively easy and additional effort investment would not further improve accuracy. For hard trials, accuracy was close to chance (floor level) and the amount of effort needed to improve accuracy may have been too much. Recent work on adaptive allocation of mental effort indeed shows that participants take into account the marginal value of effort (i.e. how much they can gain when exerting additional effort considering its cost)(Otto et al., 2022). Taking this into account, choice behavior on hard trials was likely based on risk assessment (i.e. based on the probability of being correct) rather than on careful effort-reward trade-off considerations. This is supported by our decision RT data showing that participants are slower in deciding to engage in medium effort trials compared to easy or hard trials, indicating a more deliberate choice. Future studies could reduce such differences in reward probability by individually calibrating effort levels to a participants’ own performance level like in Draper et al. (2018), thereby aiming for comparable accuracy levels across difficulty levels and comparable difficulty levels across participants. Furthermore, the current study offers important insights on the impact of inflammation on effort exertion when difficulty level is still manageable (accuracy above chance level), as compared to difficulty level approaching impossible (accuracy at or below chance level). According to the influential Motivational Intensity Theory (Brehm & Self, 1989; Silvestrini & Corradi-Dell’Acqua, 2022), effort investment is proportional to task difficulty up to a maximum limit, where difficulty is too high or the reward is not worth it anymore, leading to disengagement. Our data suggest that only below this maximum limit, mental effort exertion is affected by systemic inflammation. In sum, our results indicate that during peak inflammation levels, mental effort allocation is reduced only when the benefit of effort investment is high.

Opposite to effort, the influence of reward information on choice was not affected by systemic inflammation, indicating a selective effect of inflammation on effort. This selective effect is not attributable to changes in perception, as the LPS-induced response did not affect perceived difficulty or enjoyment ratings. This result is in line with previous animal and human endotoxemia studies, which also report changes in effort but not reward sensitivity following effort-based decision making during acute systemic inflammation (Draper et al., 2018; Lasselin et al., 2017; Yohn et al., 2016). Previous neuroimaging studies have demonstrated dissociable substrates for effort and reward processing (Hauser et al., 2017; Klein-Flugge et al., 2016; Lopez-Gamundi et al., 2021; Skvortsova et al., 2014). Specifically, effort processing seems to rely more on dorsal fronto-striatal regions such as the anterior cingulate cortex (ACC)/dorsomedial prefrontal cortex (PFC) and the putamen, whereas reward processing seems to rely more on ventral fronto-striatal regions such as the ventromedial PFC and the ventral striatum. Furthermore, neuroimaging studies that directly compared physical and mental effort-based decision making identified the dorsal ACC and dorsolateral PFC as domain-general areas (Chong et al., 2017; Hauser et al., 2017), which are regions that have also been shown to be sensitive to inflammation (Kraynak et al., 2018). Our observation that acute systemic inflammation impairs mental effort similar to what has been observed for physical effort supports the possibility that inflammation may affect these domain-general processes of effort-allocation, potentially by altering neuromodulators such as dopamine and serotonin (Cools, 2016; Kurniawan et al., 2011; Meyniel et al., 2016; Westbrook & Braver, 2015). These neuromodulators have repeatedly been shown to alter motivational processing and effort-based decision-making in both the mental (Froböse et al., 2018; Westbrook et al., 2020) and the physical domains (Chong et al., 2015; Meyniel et al., 2016; Schmidt et al., 2012). Moreover, as mental accuracy was preserved or even improved, these data speak against a direct effect of acute systemic inflammation on domain-specific implementation of effort, where plausibly effects of inflammation on cognitive and mental effort implementation would differ. This hypothesis could be further verified in future neuroimaging studies using acute immune manipulations and (physical or mental) effort-based decision-making tasks while assessing neural responses.

The current study provides evidence for a relationship between effort expenditure and fatigue, but not between effort expenditure and the extent of the inflammatory cytokine response. Neither did we observe a direct relationship between fatigue and circulating cytokine concentrations, despite our relatively large sample size. Exploration of alternative calculations of cytokine levels, such as peak IL-6 response or the AUC, or including sex as covariate also did not result in a significant relationship between fatigue, cytokines and behavioral change, strengthening our initial finding. This is in line with earlier observations that also failed to demonstrate a relationship between effort-based choice and plasma cytokine concentrations (Boyle et al., 2019; Draper et al., 2018), but is in contrast with Lasselin et al. (2020), who combined data from four different LPS studies and did report a relationship between fatigue and IL-6 concentrations across 75 participants. However, our study was not able to confirm this finding in 83 participants who all underwent the exact same study protocol. Whilst LPS administration elicits a fairly standardized biological response, fatigue has a multi-dimensional character that likely includes both inflammation-driven fatigue as well as fatigue resulting from e.g. previous effort expenditure, time of day, sleepiness, or boredom (Karshikoff et al., 2017). Similarly, effort-based choice behavior is based on not just one variable, but on input from various psychological and neural processes to optimize the desired outcome (Kurniawan et al., 2011). It might therefore be possible that the changes that we observed in decision making may capture this broader spectrum of fatigue, rather than just the physiological response to inflammation.

It needs to be noted however that the correlation between fatigue and choice behaviour was not very strong and may therefore reflect some, but not all aspects of subjective self-reported fatigue. These arguments could also explain why we did not observe a direct relationship between fatigue and cytokine concentrations. It could also explain the difference between recovery of fatigue, cytokines and body temperature at S3. Taken together, our results suggest that subjective self-reported fatigue is a better predictor for choice behavior than physiological responses to inflammation. Future investigations should investigate under which circumstances the subjective fatigue response does show a clear relation with objective measures of systemic inflammation such as the cytokine response.

## Limitations

This study has several limitations. First, learning effects across sessions are more pronounced for mental effort tasks than for physical effort tasks and difficulty levels were not adjusted to individual arithmetic capacities. Although we controlled for learning effects and accuracy by taking individual accuracy on the training trials of each session into account, it is possible that utilizing an individually tailored task may have been more sensitive. Nevertheless, the finding of reduced willingness to exert medium effort between S1-S2 despite strongly improved accuracy rates only strengthens the notion that acute inflammation also affects the willingness to exert effort. As our study design lacked a placebo condition, we cannot fully rule out out potential effects of acute inflammation on the ability to perform calculations. For example, it is still possible that LPS-induced systemic inflammation decreased the learning effect. To better control for a training effect or individual abilities, future studies could add a placebo condition, overtrain participants on the calculation task, or use mental effort tasks that are less dependent on learning.

Second, our results were not compared to a placebo condition. Therefore, we cannot state with absolute certainty that effects seen during S2 are due to the LPS-induced inflammatory response or simply reflect changes over time. To control for this, the task was assessed at three timepoints: before LPS administration, 2 hours after LPS administration (when cytokines/sickness response peak) and 5 hours after LPS administration (when cytokines/sickness response largely recovered to baseline), to show that the observed pattern of our main behavioral effect occurring during S2 is fully restored during S3. Timings of this study were similar to those in previous LPS studies (Draper et al., 2018; Lasselin et al., 2017) that did include a placebo condition and show that cytokines and sickness symptoms remain low during all three sessions after placebo. The study by Draper et al. (2018) also showed that physical efforts did not change in the placebo condition. Together, we feel confident in stating that behavioral changes during the study are most likely ascribed to the LPS-induced inflammatory response.

## Conclusion

In conclusion, this is the first study to investigate relationships between mental effort-based decision making, fatigue and systemic inflammation in a large sample. Our results show that the LPS-induced inflammatory response in young healthy adults reduces the willingness to invest mental effort, without affecting reward sensitivity. This change in decision making occurred despite increased mental task accuracy, suggesting a reorganization of motivational priorities rather than an inability to exert mental effort, similar to what has been observed for physical effort-based decision making. These changes in mental effort allocation were associated with changes in subjective fatigue, but not with objective parameters of systemic inflammation.

Our results indicate that inflammation may not necessarily affect domain-specific cognitive functioning, but could also affect domain-general effort-allocation processes. While acute effects of inflammation might not be directly translatable to long-term effects, understanding the origin and mechanisms behind inflammation-induced acute fatigue is an essential first step to understanding mental fatigue and cognitive problems in inflammation-associated conditions. To develop a full picture of fatigue-related motivational changes, future research in fatigued patient populations should not only measure general cognitive functioning but also investigate to what extent inflammatory effects on domain-general motivational processes may contribute to mental fatigue problems. This could provide essential insights for the development of new treatments targets such as effort perception or effort-allocation processes, progressing from symptomatic to curative treatment of fatigue and cognitive symptoms associated with illness and medical treatment.

## Acknowledgements

We would like to acknowledge all interns and members of the intensive care research department of the Radboud University Medical Center for the opportunity to join this study and for all their practical help, particularly Niklas Bruse, Nicole Waalders, Jelle Gerritsen and Corisjo Schroevers.

