## Supplementary materials for "Fatigue during acute systemic inflammation is associated with reduced mental effort expenditure while task accuracy is preserved"

**Inclusion criteria**

- Written informed consent
- Age ≥18 and ≤35 years
- Healthy (as confirmed by medical history, physical examination, electrocardiography, laboratory tests)

**Exclusion criteria**

- Pregnancy (confirmed by negative result of pregnancy tests prior to endotoxin challenge)
- Use of any medication
- Smoking
- History or signs of atopic syndrome (asthma, rhinitis with medication and/or eczema)
- Known anaphylaxis or hypersensitivity to the non-investigational products or their excipients.
- History or signs of hematological disease:
    - Thrombocytopenia (<150*10^9/ml) or anemia (hemoglobin < 8.0 mmol/L)
    - Abnormalities in leukocyte differential counts
- History, signs or symptoms of cardiovascular disease, in particular:
    - Previous spontaneous vagal collapse
    - History of atrial or ventricular arrhythmia
    - Cardiac conduction abnormalities on the ECG consisting of a 2nd degree atrioventricular block or a complete left bundle branch block
    - Hypertension (defined as RR systolic > 160 or RR diastolic > 90)
    - Hypotension (defined as RR systolic < 100 or RR diastolic < 50)
- Renal impairment (defined as plasma creatinine >120 μmol/l)
- Liver enzyme abnormalities (above 2x the upper limit of normal)
- Medical history of any disease associated with immune deficiency
- Signs of infection (CRP > 20 mg/L, WBC > $12x10^9$/L or < 4 $x10^9$/L)
- Clinically significant acute illness, including infections, within 1 month of the first endotoxin challenge

- Previous (participation in a study with) endotoxin (LPS) administration
- Any vaccination within 3 months within of the first endotoxin challenge
- Participation in a drug trial or donation of blood within 3 months prior to first endotoxin challenge
- Recent hospital admission or surgery with general anesthesia within 3 months prior to first endotoxin challenge
- Use of recreational drugs within 1 month of the first endotoxin challenge
- Inability to personally provide written informed consent (e.g. for linguistic or mental reasons) and/or take part in the study
- Unwillingness to be informed about potential chance findings

**Table S1.** Inclusion and exclusion criteria

One hour before LPS administration:

- Oral assessment of physical sickness symptoms on a 5-point Likert scale (headache, muscle pain, back pain, nausea, shivers, vomiting) during past week
- Multidimensional Fatigue Index (MFI), during past week
- Expected sickness response (one question: “I think that, today, I will feel [blank] what I am used to when I am sick”, with a VAS scale from “much worse than - much better than”)

After LPS administration:

- Oral assessment of physical sickness symptoms on a 5-point Likert scale (headache, muscle pain, back pain, nausea, shivers) (every 30 minutes until 360 minutes post-LPS)
- Profile of Moods State Questionnaires (POMS) (every 60 minutes until 360 minutes post-LPS)
- Positive and Negative affect scale (PANAS) (every 60 minutes until 360 minutes post-LPS)
- State Anxiety Inventory (STAI) (every 60 minutes until 360 minutes post-LPS)
- Acute Multifactorial Fatigue Inventory (aMFI) (every 60 minutes until 360 minutes post-LPS)

**Table S2.** Overview of all questionnaires during testing day

| *Condition* | | *Session* | | |
|---|---|---|---|---|
| Effort level | Reward level | 1 | 2 | 3 |
| Easy | Low | 93.6(2.7) | 94.2(2.5) | 95.9(2.2) |
| Easy | Middle | 97.5(1.7) | 98.4(1.4) | 99.3(0.9) |
| Easy | High | 99.2(1.0) | 99.8(0.4) | 99.6(0.7) |
| Medium | Low | 61.5(5.3) | 62.9(5.2) | 70.5(4.9) |
| Medium | Middle | 82.9(4.1) | 79.9(4.3) | 85.9(3.8) |
| Medium | High | 91.2(3.1) | 84.5(3.9) | 92.2(2.9) |
| Hard | Low | 23.3(4.6) | 32.0(5.1) | 41.1(5.3) |
| Hard | Middle | 44.8(5.4) | 54.8(5.4) | 61.1(5.3) |
| Hard | High | 64.6(5.2) | 68.7(5.0) | 74.2(4.8) |

**Table S3:** mean (SEM) of % accepted offers for each combination of effort and reward level per session

| | *Session* | | |
| --- | --- | --- | --- |
| Effort level | 1 | 2 | 3 |
| Easy | 80.0(1.6) | 84.7(1.6) | 85.3(1.6) |
| Medium | 65.8(2.1) | 75.3(1.7) | 72.2(1.9) |
| Hard | 58.2(2.3) | 64.6(2.4) | 70.6(2.1) |

**Table S4:** mean (SEM) of % correct calculations for each effort level per session during training phase

| | | Df | Chisq | p |
|---|---|---|---|---|
| *Main effects* | Session | 2 | 4.981 | 0.083 |
| | Reward | 2 | 133.707 | <0.001 |
| | Effort | 2 | 263.157 | <0.001 |
| *Effects of interest* | Effort*Session | 4 | 17.020 | 0.002 |
| | Reward*Session | 4 | 2.502 | 0.644 |
| | Effort*Reward*Session | 8 | 13.554 | 0.094 |

**Table S5.** Results from generalized estimating equation (GEE) analysis (N=85)

| | | S1-S2 | | | S2-S3 | | | S1-S3 | | |
|---|---|---|---|---|---|---|---|---|---|---|
| | | Df | Chisq | p | Df | Chisq | p | Df | Chisq | p |
| *Main effects* | Session | 1 | 0.029 | 0.866 | 1 | 4.650 | 0.031 | 1 | 1.098 | 0.295 |
| | Reward | 2 | 147.084 | <0.001 | 2 | 150.804 | <0.001 | 2 | 248.308 | <0.001 |
| | Effort | 2 | 206.030 | <0.001 | 2 | 72.594 | <0.001 | 2 | 138.922 | <0.001 |
| *Effects of interest* | Effort*Session | 2 | 14.771 | <0.001 | 2 | 9.049 | 0.011 | 2 | 1.971 | 0.373 |
| | Easy | 1 | 1.807 | 0.179 | 1 | 0.081 | 0.776 | - | - | - |
| | Medium | 1 | 9.246 | 0.002 | 1 | 17.607 | <0.001 | - | - | - |
| | Hard | 1 | 0.305 | 0.581 | 1 | 0.1182 | 0.731 | - | - | - |
| | Reward*Session | 2 | 0.200 | 0.905 | 2 | 2.432 | 0.296 | 2 | 0.794 | 0.673 |
| | Effort*Reward*Session | 4 | 8.801 | 0.066 | 4 | 8.2939 | 0.081 | 4 | 3.1847 | 0.5274 |

**Table S6.** Results from generalized estimating equation (GEE) analysis (N=85), session comparisons

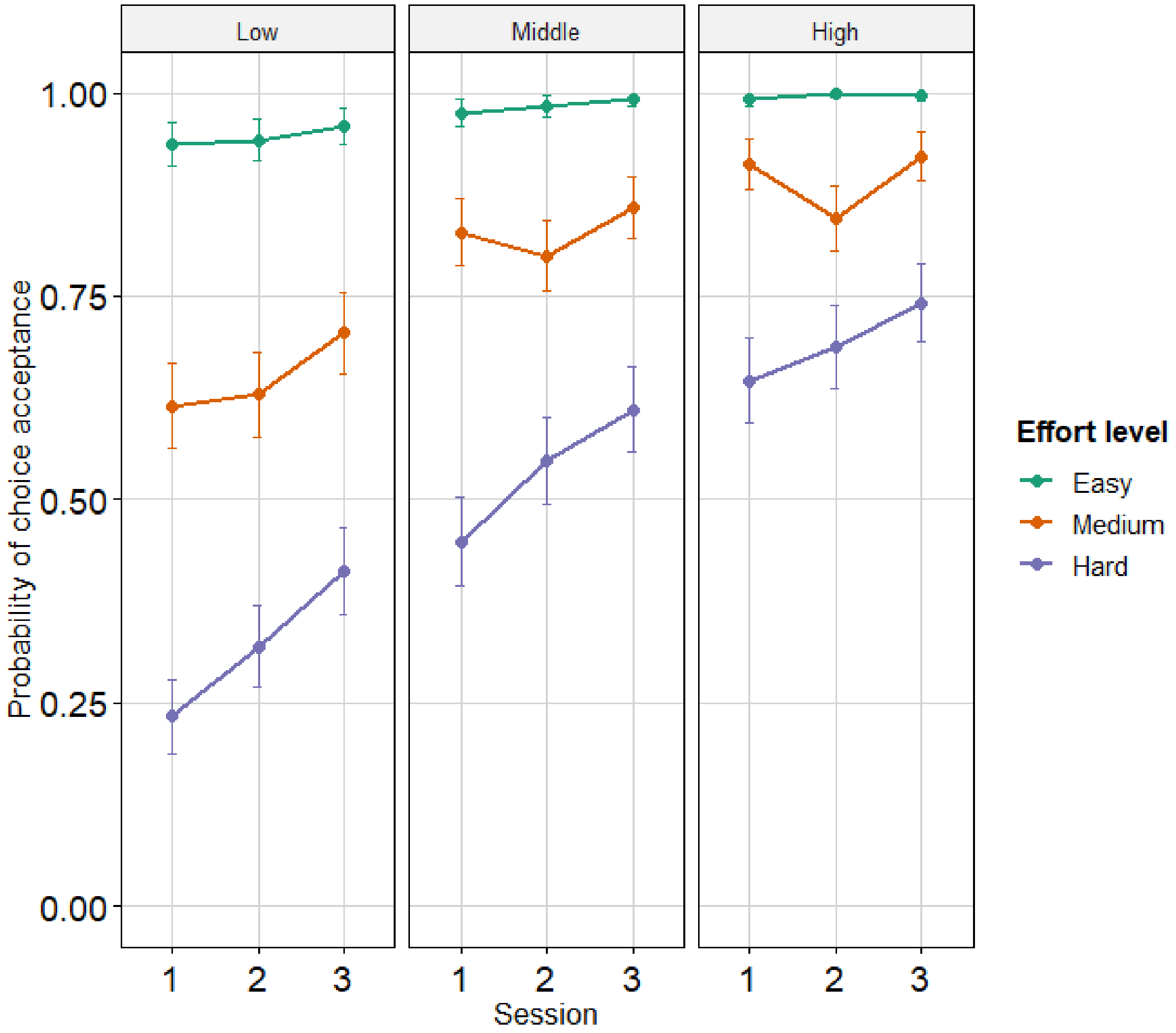


**Figure S1.** Raw uncorrected means for choice acceptance split for effort and reward level for all three sessions

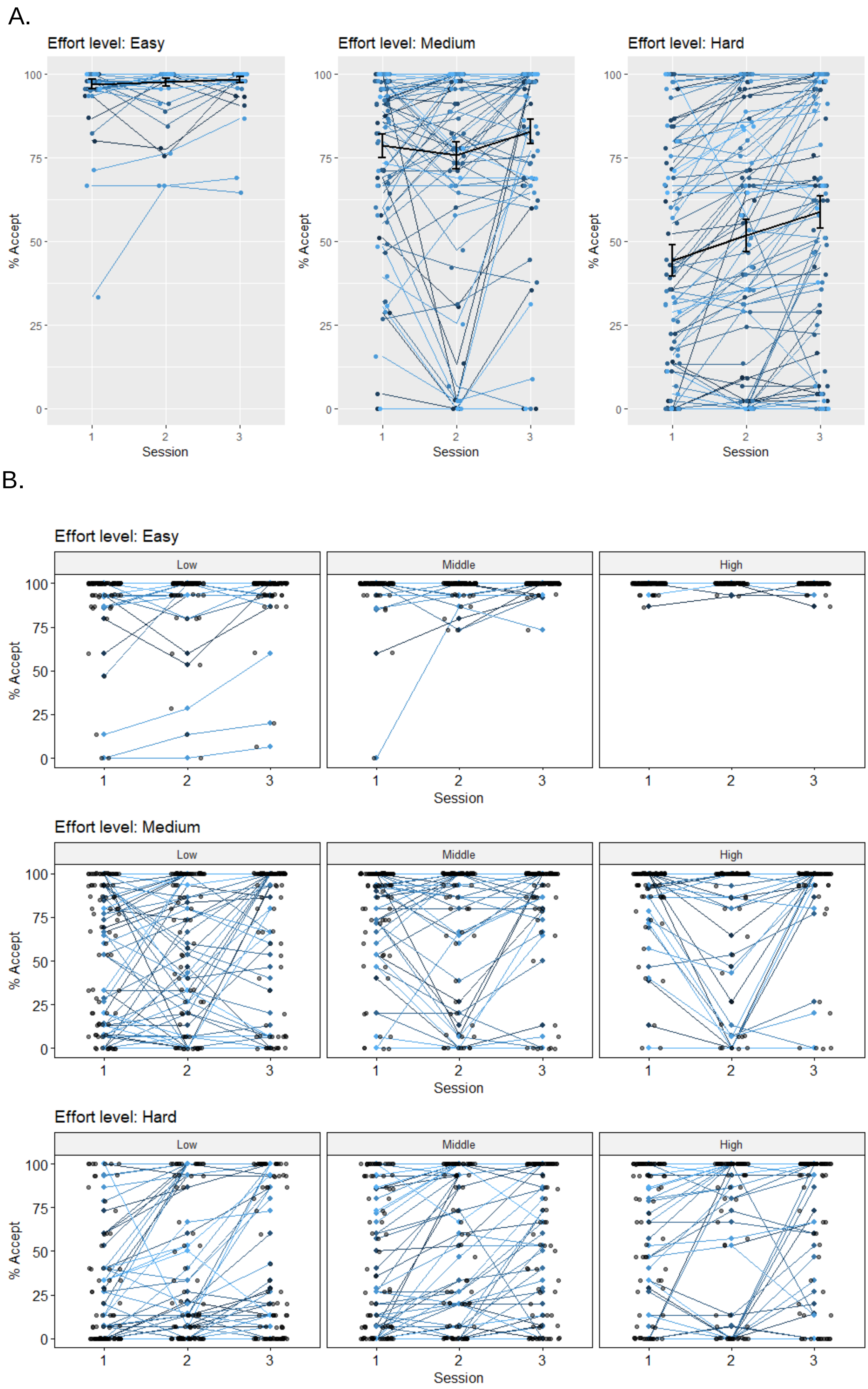
A.
Effort level: Easy
Effort level: Medium
Effort level: Hard
% Accept
Session
B.
Effort level: Easy
Effort level: Medium
Effort level: Hard
Low
Middle
High
% Accept
Session

Figure S2: Individual trajectories and means of the raw uncorrected acceptance data for all three sessions. A) split for effort level, and averaged across reward level. Different colors represent different subjects.B) split for effort level (top=easy, middle = medium, bottom = hard) and reward level (left = low, middle = middle and right = high). Different colors represent different subjects.

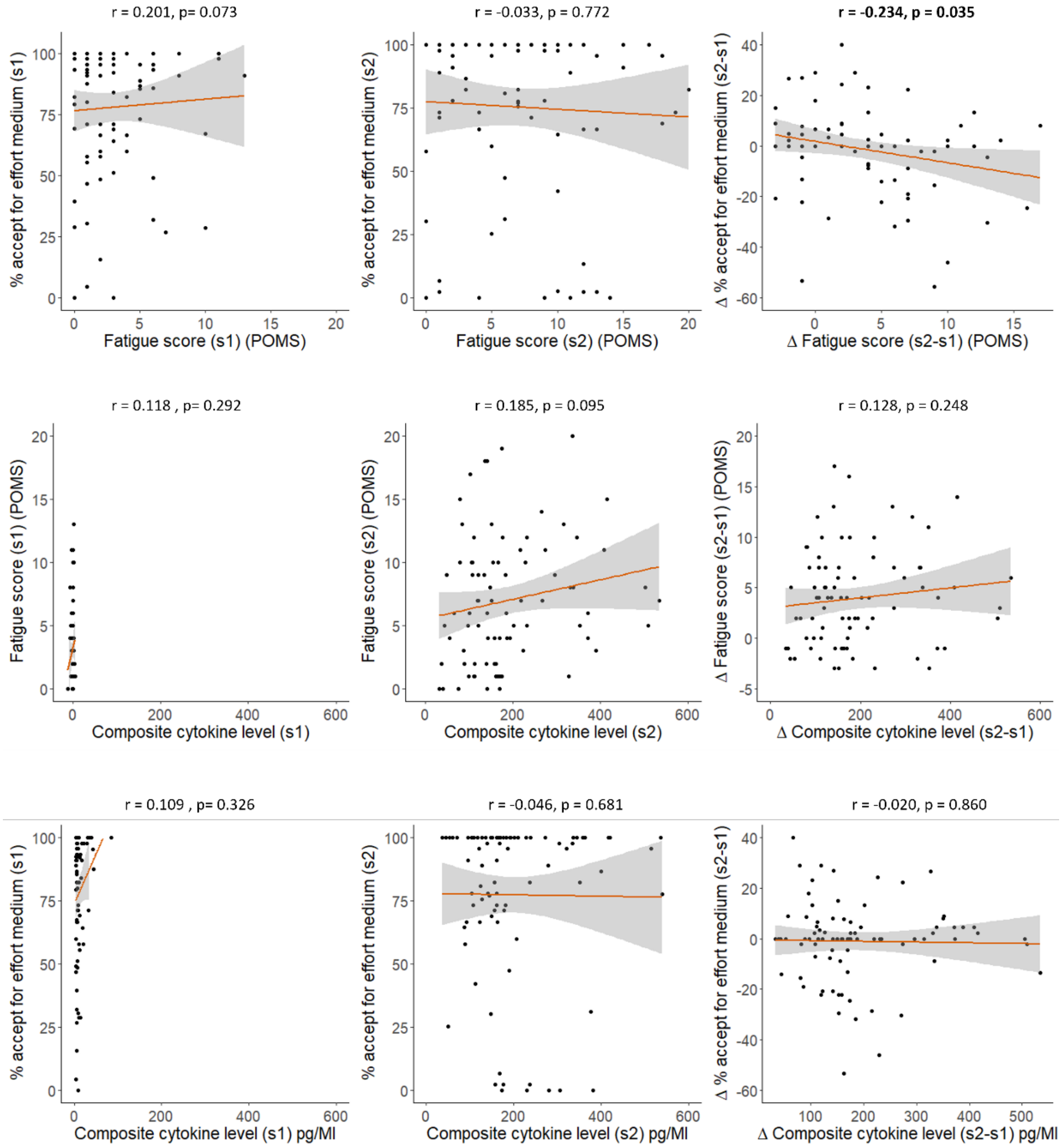


FigureS3: Correlations between acceptance and fatigue (upper row), fatigue and cytokines (middle row) and Accept and cytokines (lower row for session 1(left column), session 2 (middle column) and the difference between session 2 and session1 (right column) for the medium effort trials. Correlations that include acceptance rates are corrected for accuracy.

Figure S4: Point structure of the task in relationship to accuracy.
When an offer is accepted subjects receive the low, medium or high reward when correct and lose 1 point when incorrect. When an offer is rejected, subjects always receive 4 points. In blue, the expected reward is shown given the accuracy level on the left. By using this structure, acceptance of the offer is only beneficial in terms of expected rewards, when accuracy exceeds 0.62 for the highest rewards (i.e. middle and dark blue areas). When employing an effortless guessing strategy, rejection is the better option.

| | | | | | |
|---|---|---|---|---|---|
| reject = better | | | | | |
| indifference point | | | | | |
| accept = better | | | | | |
| | **Reward level:** | **low** | **medium** | **high** | |
| | Reject | Accept | Accept | Accept | |
| | 4 | -1 | -1 | -1 | |
| Accuracy level | 4 | 5 | 6 | 7 | Better option, given accuracy and expected reward |
| 0,3 | 4 | 0,8 | 1,1 | 1,4 | reject everything |
| 0,4 | 4 | 1,4 | 1,8 | 2,2 | reject everything |
| 0,5 | 4 | 2 | 2,5 | 3 | reject everything |
| 0,6 | 4 | 2,6 | 3,2 | 3,8 | reject low rewards |
| 0,7 | 4 | 3,2 | 3,9 | 4,6 | reject lowest reward |
| 0,8 | 4 | 3,8 | 4,6 | 5,4 | accept highest rewards |
| 0,9 | 4 | 4,4 | 5,3 | 6,2 | accept everything |
| 1 | 4 | 5 | 6 | 7 | accept everything |
